# Differences in the spatial and temporal patterns of head motion during MRI of adults and infants

**DOI:** 10.1101/114447

**Authors:** Rhodri Cusack, Annika C. Linke, Leire Zubiaurre-Elorza, Hester Duffy, Charlotte Herzmann, Bobby Stojanoski, Victor K. Han, David S.C. Lee, Conor Wild

**Author notes:** Corresponding author:* Rhodri Cusack, Brain and Mind Institute (NSC 203), University of Western Ontario, London ON N6A 5B7, Canada.

## Abstract

**Aim:** Head motion has a profound effect on MRI, and will contaminate comparisons of function or structure between groups that move differently. This work compares adults and infants. Infants might move differently for physical, physiological and cognitive reasons, but so far these differences have not been quantified.

**Methods:** The spatial modes and total magnitude of motion in the MRI scanner were measured (N=211). The effects of group (infant vs. adult) and stimulation paradigm (auditory vs. visual) were evaluated.

**Results:** Spatial modes of motion were found to be distinct between infant and adult groups. Infants had less anterior-posterior translational motion, but greater motion in other dimensions, often with complex multi-axis patterns. In magnitude distribution, sleeping infants often remained more still than adults, but when movement did occur it was more extreme and abrupt. Two groups of adults presented with different stimulation showed similar shapes of motion.

**Conclusion:** The spatial modes and magnitude distribution of motion differed substantially between groups, and must be considered carefully as a confound in comparisons of structure or function. The abruptness and magnitude of movement suggests that for infants relative to adults post-processing strategies such as de-noising are likely to be more effective than prospective motion correction.

**Key notes:**

- Quantified the spatial and temporal distribution of motion during MRI in 211 adults and neonates
- The different spatial modes in adults and infants were visualized and statistically contrasted
- The magnitude of motion had “heavier tails” in infants, with more still periods, and more large movements, than adults.

---

Participant motion is a serious problem in MRI of the head, affecting acquisitions that measure function, anatomy and connective tracts. For example, in functional connectivity MRI (fc-MRI), two brain regions are determined to be part of the same network if they show a similar pattern of fluctuation in activity over time. Noise introduced by motion can strengthen or weaken these correlations, and may have a structured spatial pattern: the motion-induced signal is often similar for regions that are close together and dissimilar for regions far apart, increasing the relative strength of short-rather than long-range connections [1,2]. As motion may also differ between groups under study (e.g., children are more likely to move than adults), great care must be taken in interpreting a difference between groups in any measure that is affected by motion [3].

Motion is a particular problem for some populations. For example, older adults may have kyphosis of the spine and find it uncomfortable to lie supine for an hour or more. Also, some subjects - such as patients in a vegetative state, pre-linguistic infants, or animal models - cannot understand the need to remain still during a scan. As a result, motion tends to be a significant barrier to successfully scanning these groups. Sometimes it is possible to modify MRI protocols to mitigate the effects of motion. One simple approach is to use sequences that are more robust, such as acquiring rapid 2D slices rather than slower 3D volumes. A promising active area of research is on prospective motion correction MRI sequences which measure motion with fast “navigator” acquisition [4,5], by analysis of the incoming stream of MRI data [6,7], or through optical [8,9] or ultrasonic [10] tracking. The MRI acquisition is then corrected prospectively. This field is not without a number of challenges [11]. None of these methods is fully effective in preventing motion artifacts, and several come with significant costs in terms of acquisition time or resolution. Also, due to the technical requirements of these methods (e.g., custom equipment, scanner interfaces, and pulse sequences) they are not readily available at many scanning sites, including the clinical settings where only some patients groups, such as premature infants, can be scanned. Nonetheless, prospective motion correction tools offer exciting potential, particularly for patient groups where motion is unavoidable.

The effect of motion can also be mitigated at the time of data analysis with retrospective correction methods. Most common of these is rigid-body realignment, in which each 3D volume is realigned to a reference image, and estimates of movement (three rotations, three translations) are regressed out of the fMRI time series. However, motion-induced artifacts remain in the data even after this process and for scientific or clinical research that compares groups, it is common to exclude participants that move more than some predetermined threshold from further analysis. This comes with substantial costs - like many, our laboratory typically rejects around 10-20% of healthy young adult volunteers during fMRI [12-17]. In infant studies, rejection rates are much higher [18-20]. Rejection itself may also introduce systematic artifacts, as found recently in a study of multiple sclerosis, where participants that were rejected due to motion were found to have a different cognitive profile than those left in the analysis [21]. In addition to rejection, other techniques that aim to remove noise during analysis are based on independent component analysis [22-24], and the modeling out of estimates of nuisance variables from the time series [25].

This plethora of potential acquisition and analysis strategies often begs the question, which are necessary and will perform best for any given study? The answer to this question will depend on the form of the movement in the participants to be studied. Here, as a proof of principal we characterize motion in a group in which it presents a particular challenge: neonates. We compare the spatial and temporal form of motion in neonates with that of healthy young adults. There are reasons to expect that adults and neonates may move in different ways. Neonates have poorly developed musculature in the neck, and find a nodding action difficult. They are smaller allowing them more freedom of movement in the scanner, but were swaddled during scanning, so don’t have freedom to move their arms and legs. To visualize the modes of motion, we performed principal components analysis of the motion trajectories across scans in the adult and infant groups. We hypothesized that motion would be different in extent, in its spatial modes, and its distribution through time.

## Methods

The data presented here come from fMRI, but the images were used here solely to measure the motion of the head during the scanning session.

### Participants

In a first adult group, 40 scanning sessions were acquired using a Siemens Tim Trio 3T MRI scanner. 23 participants were recruited (age 25.4+/-4.6 years; 9 male, 14 female) and 17 participated in two sessions. Rapid snapshots of brain position were acquired using a highly accelerated gradient-echo EPI sequence (Center for Magnetic Resonance Research, University of Minnesota) with multiband acceleration factor 3 and GRAPPA iPat acceleration of 2. 32 slices were acquired with a matrix size of 70x70 and a voxel size of 3 x 3 x 3 mm (not inclusive of a 10% slice gap), TE=25 ms, and TR=850 ms. Between 1 and 3 series were available from each scanning session (2.75 +/-0.49, 110 series total). We selected the first 7 minutes 2 seconds of data (422 volumes) from each scanning series, to match the duration of the infant acquisitions. Participants performed a visual short-term memory task, not relevant to the current project.

In infants, 28 scanning sessions were conducted on a GE MR450W 1.5T scanner. Neonates were swaddled and snuggled into a MedVac Immobilisation Blanket. They were monitored with a noise-cancelling microphone (Optoacoustics), and pulse oximetry. Many appeared to sleep through the scan. These research EPI scans were added to clinical scans at Children’s Hospital, London Health Sciences Centre, London Ontario. Infants were inpatients of the Neonatal Intensive Care Unit (NICU) and were either born very prematurely (<29 weeks), small for gestational age, or had sustained some other event that placed them at higher risk of neurological injury (e.g., suspected hypoxia). Age at scan was 38 +/-2.5 weeks post menstrual age (data from two infants unavailable) and their age at birth 32 +/- 6 weeks (data from one infant unavailable). 21 were male and seven were female. EPI data were acquired using a GE product gradient-echo sequence with 22 slices, matrix size 44x44, voxel size 3x.3x3mm (not inclusive of a 25% slice gap). Data were zero-padded by the scanner during reconstruction to 64x64. An echo time (TE=60 ms) longer than is typically used for adults was used to compensate for the higher T2* in infants [26]. The TR was 1970ms and 220 volumes were acquired in each series (7 mins and 2 secs). Between 1 and 4 series were acquired per infant (3.0 +/- 1.1, 85 series total). Auditory stimuli were presented, which are not relevant to the current project.

A possible criticism of comparing movement between the first adult group and the infants is that although the stimulation was irrelevant to our analysis, perhaps the presence of a visual stimulus in adults affected their patterns of movement. We therefore analyzed data from a smaller second adult group that were listening to exactly the same stimuli as the infants. MRI was acquired on a Siemens Prisma 3T scanner at the Centre for Functional and Metabolic Mapping, in 16 adults (age 23.3 +/- 5.0 years; 11 female, 5 men) using a multiband gradient-echo EPI sequence (Center for Magnetic Resonance Research, University of Minnesota) with multiband acceleration factor 4 and no GRAPPA iPat acceleration. 36 slices were acquired with a matrix size of 64x64 and a voxel size of 3 x 3 x 3 mm (not inclusive of a 10% slice gap), TE=30 ms, and TR=686 ms. Three sessions each of 610 volumes (6 mins 58 secs) were acquired.

### Analysis

#### The automatic analysis system, version 4

(www.github.com/rhodricusack/automaticanalysis) was used to pipeline analyses with SPM 8 (Wellcome Trust Centre for Neuroimaging, London, UK). Data were converted to Nifti format and realigned to the first image of the time series using SPM. The six motion parameters, describing translation (*x* - left/right, *y* - anterior/posterior and *z* - superior/inferior) and rotation (*pitch* - chin up/down, *roll* - top of head left/right, and *yaw*-nose left/right) were then extracted for each scan and processed using custom code written in Matlab (Mathworks, Natick, Massachusetts).

### Spatial modes of movement

To visualize whether adults and infants have different ways of moving, we performed principal component analysis to reduce the six time series of motion for each group into spatial modes, concatenating timeseries for subjects and sessions within each group. We retained only those components (i.e., modes) that explained > 80% of the variance in the motion trajectories. These were shown in a bar graph, but to further assist in the visualization of the modes of movement, we rendered the motion in a three dimensional model. MakeHuman 1.02 (http://www.makehuman.org) was used to obtain meshes of the adult and infant, which were then cropped to just the head using a boolean modifier in Blender 2.72 (http://www.blender.org), and the motion modes from the principal components rendered using Matlab 2014b (http://www.mathworks.com).

### Quantifying movement

We calculated the covariance matrix for the 3 translations (in *mm*) and 3 rotations (in *degrees*) for each participant, concatenating series where more than one was available. Each point in this 6x6 covariance matrix was contrasted between infants and adults using heteroscedastic two-sample t-tests. This comparison was done in two ways. First, the raw covariance values were compared. This allowed statistical assessment of how the total amount of movement (e.g., variance in *x* position), and the relationship between different axes of movement (e.g., covariance between *x* position and *y* position), differed across groups. Second, we wanted to assess whether, independently of any difference in magnitude, the spatial pattern of movement differed between groups. To do this, the covariance matrix for each subject was normalized by the total single axis movement (i.e., by the *trace* of the covariance matrix). The elements in these normalized covariance matrices were then compared across groups. We compared the infant group (N=28) to a group of adults (group 1, N=40) and a smaller group of adults listening to the same stimulation as the infants (group 2, N=16).

### Magnitude of movement

To provide a measure of the magnitude of the movement that was independent of the spatial mode, a summary statistic was calculated that combined the possible translational and rotational modes in a principled way - the *root mean square* displacement of each voxel within a sphere [27] that approximates the size of the brain. For adults, this was taken to be the value used by Jenkinson (r=8 cm). For infants at birth, we scaled this value, approximating it as following head circumference, using an adult male head circumference of 57 cm and a newborn head circumference of 34.5 cm (WHO, http://www.who.int/childgrowth/standards/hcforage/en), giving r=4.84cm. Three measures were calculated. The absolute movement through the scan was the RMS displacement relative to the first volume. The relative movement was the RMS displacement of adjacent scans. However, the adult and infant MRI sequences had different TRs (adult group 1 – 850, group 2 – 686, infant - 1970 ms) and so the RMS difference between adjacent scans may relatively underestimate motion in the adults. To account for this, we also calculated the RMS displacement across matched intervals (adult group 1 – 7 TRs, 5.95 s; group 2 – 9 TRs, 6.17 s; infant - 3 TRs, 5.91 s). To the small extent (1-5%) that these intervals are still mismatched across groups, the direction of the mismatch is opposite to the difference between adjacent scans. These per-scan RMS measures were then summarized across all of the scans in all of the sessions of each subject. Random-effects statistics were then calculated across subjects. We quantified the RMS displacement in three ways: the mean across scans; the number of very small movements (<0.1mm); and the number of large movements greater than the size of a voxel (>3mm).

## Results

### Example trajectories

Sixteen randomly chosen and representative examples of motion in adults (from group 1) and infants are shown in Figure 1. It can be seen by eye that infants show different distributions and temporal profiles of movement compared to adults. Some infants remain very still throughout the scan, but many show high-amplitude spikes of movement.

**Figure 1.**
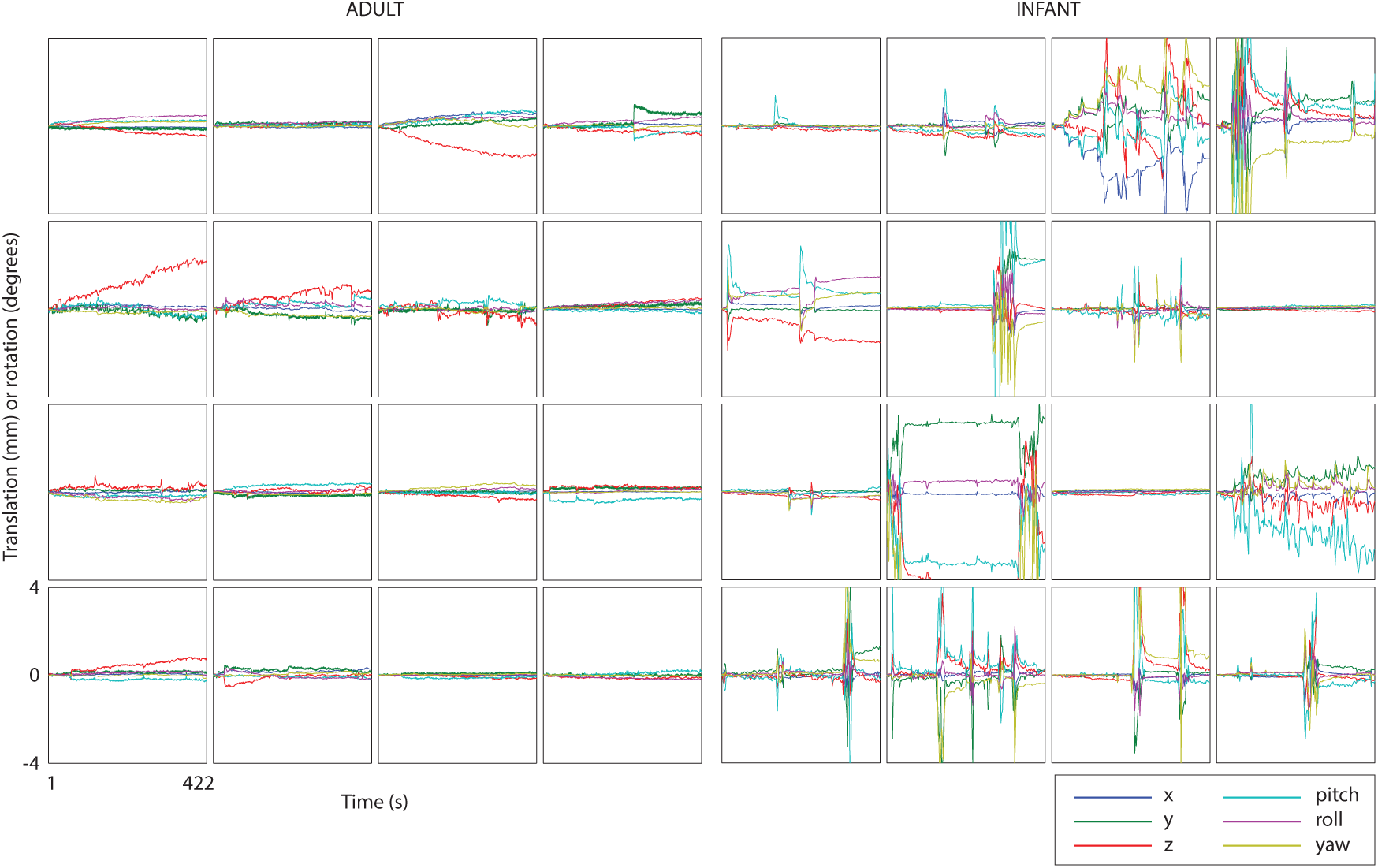
Sixteen representative motion tracks from adult group 1 and the infant group. The six curves in each plot denote x, y, z translation (in mm) or rotation (in degrees) around the x, y and z axes (pitch, roll, yaw respectively). The axis ranges are the same for all plots.

### Spatial modes of movement

The main spatial modes of movement were quantified using principal component analysis. It was found that in adults (both groups 1 and 2) and neonates, three principal components accounted for more than 80% of the variance. Figure 2 shows these first three components for the three groups. The adult patterns are reassuringly similar, although the first two principal components (which were of similar magnitude) were reversed in order in group 2 relative to group 1. The infant pattern appeared different to both adult groups, with adults showing movements weighted towards a single axes of movement (notably z translation, pitch and roll) but infants showing a more complex wriggle with complex patterns of interrelated movements along multiple axes.

**Figure 2.**
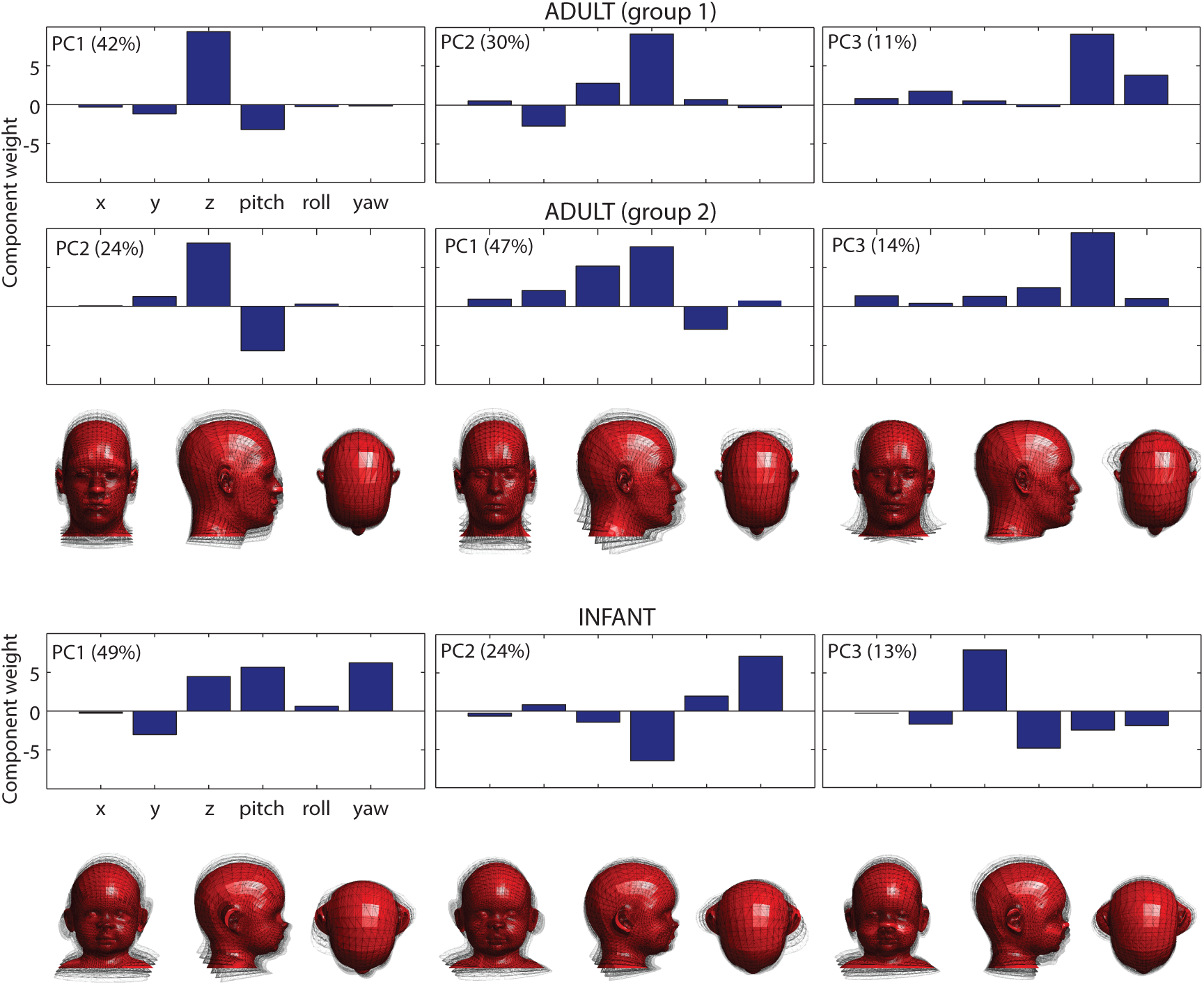
Key modes of motion for the adults (groups 1 and 2) and infants, derived using principal components analysis. The top three components for each group are shown, and the percentages in brackets shows the variance explained by each component. Note that components 1 and 2 are presented in reverse order for adult group 2, so that they correspond to group 1. The three dimensional renderings visualize the three modes of motion for the adults (group 1) and the infants. The axis ranges are the same for all plots.

### Quantifying movement

Figures 3a-c show the mean normalized covariance matrices for the infant and two adult groups. Values along the leading diagonal of each covariance matrix show the variance of motion for the three translations and three rotations. Statistical comparison of the raw covariance values showed that infants move more than the first larger adult group (Figure 3d) in all ways except yaw. Similarly, compared to the second adult group (Figure 3e), the infants showed more motion in all ways except yaw and z translation. There was also greater covariance of many pairs of axes in infants, but from these raw covariances it is difficult to determine whether this is merely a reflection of greater single-axis motion.

**Figure 3.**
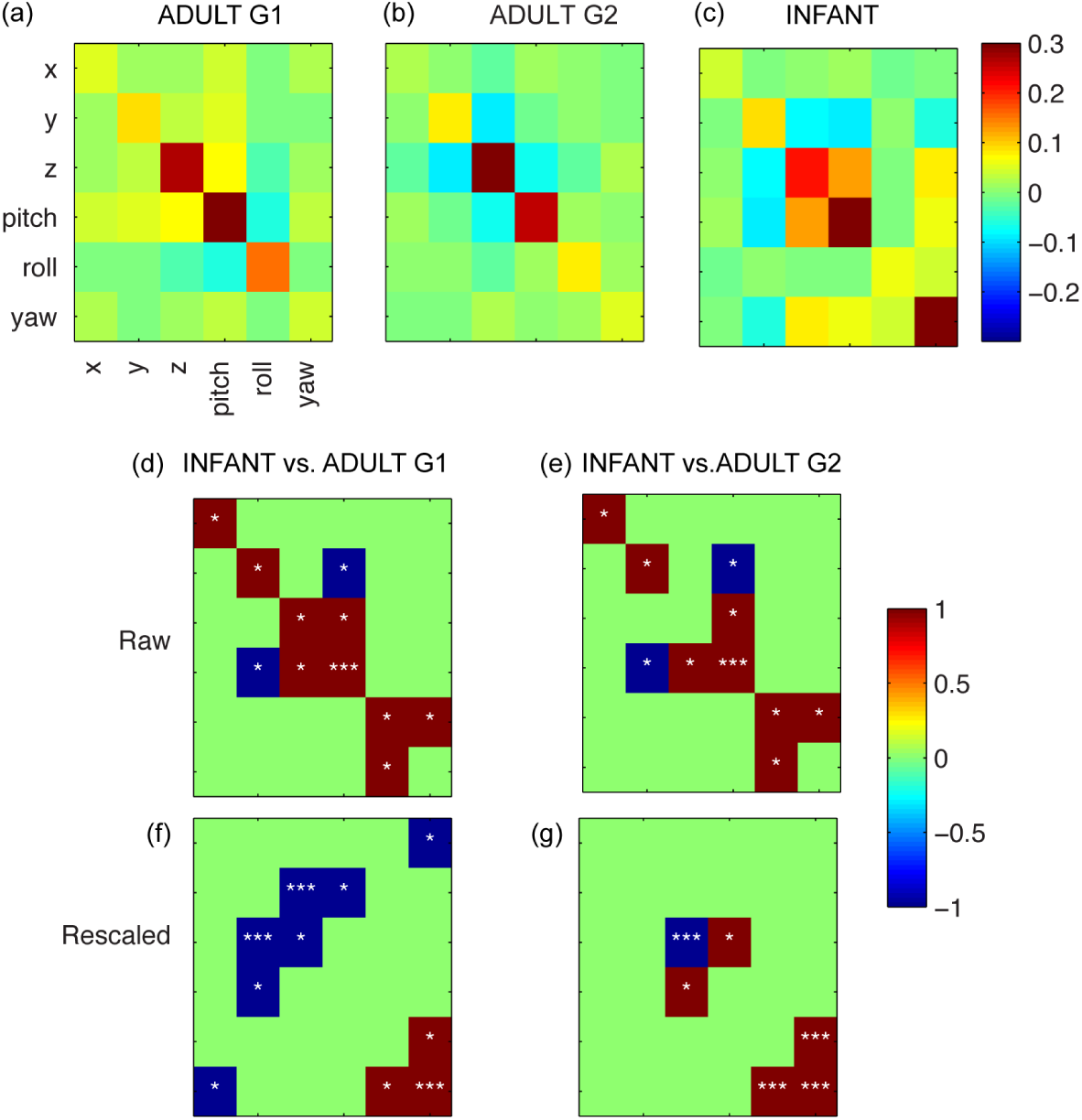
To test how modes of motion were statistically different, we examined the covariance matrices of the six motion parameters. Along the leading diagonal, the covariance matrix captures the magnitude of motion on individual axes, and off this diagonal, the relationship between different kinds of motion. (a-c) show the normalized covariance matrices for each group, averaged across subjects. (d) and e) show the statistical comparison of the infants to each of the adult groups, using the raw data, which captures the magnitude as well as pattern of motion (red infants>adults, blue adults>infants, green not significant; * p<0.05; ** p<0.01; *** p<0.001). (f) and (g) shows the statistical comparison of the normalized data, which captures the pattern of motion with the overall magnitude (the trace) scaled out. The condition ordering is the same for all plots.

Comparison of the normalized covariance matrices allowed a test of whether particular axes showed more movements than others in the different groups, or whether pairs of axes were more related. The patterns were quite different. Relative to the total amount of motion, there was less translation along the *z*-axis in infants, but more yaw, and a greater correspondence between *z* axis motion and pitch, and roll with yaw. In adults, *z* translation (along the long axis of the body) probably reflects movement of the legs or torso, a type of motion not available to the swaddled infants. However, the infants can move their heads side to side, to create yaw. Many off-diagonal elements were also significant, suggesting that the coupling between movements along different axes is quite different in adults and infants.

### Magnitude of movement

To assess the magnitude of movement independently of the spatial mode, an RMS summary was calculated that combines translations and rotations in a principled way, and accounts for head size. Figure 4 and Table 1 show the results. One key result is that when motion is measured relative to the first scan, infants showed a similar or lower degree of movement than both groups of adults: no statistical difference in the mean RMS or in the proportion of scans with large movements (>3 mm), and a greater number of scans with very small shifts (<0.1 mm). However, greater infant motion was found on a faster timescale. When RMS movements were measured between adjacent scans, or across a time interval matched between groups, there was greater prevalence at the extremities, with more small (<0.1mm, compared to adult group 2 only) and large (>3mm, both groups) movements. This rapid motion in infants is likely to be quite pernicious to fMRI analyses, as it will not be removed by high-pass filtering, which is standard in fMRI pipelines and will remove much of the gradual drift seen in adults.

**Figure 4.**
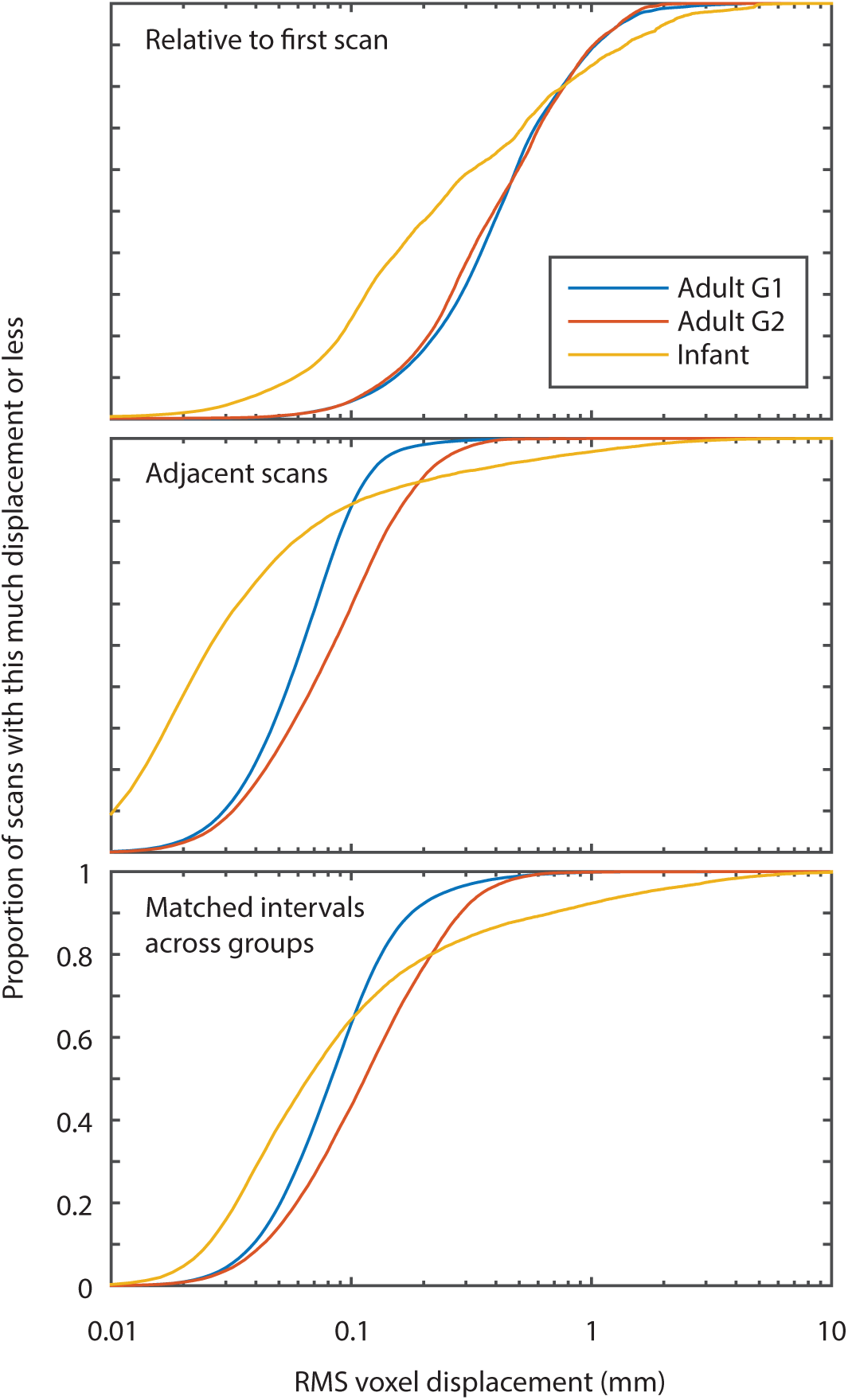
The cumulative distribution of movement magnitudes, displayed as the proportion of scans with RMS displacement less than or equal to the value on the x-axis. The top panel shows this relative to the first scan; the middle panel for adjacent scans; and the bottom panel for a time interval ~ 6 s. The infants are more extreme, with more scans with very small movements (yellow curve above blue and red on left of plot), and more with very large movements (yellow curve below blue and red on right of plot). Table 1 shows accompanying statistics. The axis ranges are the same for all plots.

**Table 1.** Quantifying the magnitude of motion in each of the groups through the root-mean square (RMS) displacement of the voxels across the brain. “Mean” shows the mean of this RMS across scans, “<0.1mm” the proportion of scans where the RMS was less than 0.1mm, and “>3mm” the proportion of scans where the RMS was greater than 3mm. Relative to adults, infants showed greater extremities of motion - they were sometimes very still, and sometimes moved a lot.

|  | Adult G1 | Adult G2 | Infant | Infant vs. Adult G1 |  | Infant vs. Adult G2 |  |
| --- | --- | --- | --- | --- | --- | --- | --- |
| *Relative to first scan* |  |  |  |  |  |  |  |
| Mean | 0.53+/-0.27 | 0.51+/-0.21 | 0.49+/-0.35 | t(49)=-0.44 | NS | t(42)=-0.16 | NS |
| <0.1mm | 0.04+/-0.04 | 0.04+/-0.03 | 0.25+/-0.18 | t(29)=5.96 | *** | t(29)=5.91 | *** |
| >3mm | 0.00+/-0.02 | 0.00+/-0.00 | 0.02+/-0.05 | t(36)=1.24 | NS | t(27)=1.76 | NS |
| *Adjacent scans* |  |  |  |  |  |  |  |
| Mean | 0.11+/-0.04 | 0.15+/-0.06 | 0.30+/-0.20 | t(29)=5.30 | *** | t(34)=4.01 | *** |
| <0.1mm | 0.63+/-0.20 | 0.43+/-0.23 | 0.64+/-0.18 | t(61)=0.34 | NS | t(26)=3.22 | ** |
| >3mm | 0.00+/-0.00 | 0.00+/-0.00 | 0.02+/-0.02 | t(27)=4.61 | *** | t(27)=4.62 | *** |
| *Matched time interval* |  |  |  |  |  |  |  |
| Mean | 0.07+/-0.02 | 0.10+/-0.03 | 0.12+/-0.08 | t(28)=3.08 | ** | t(40)=0.92 | NS |
| <0.1mm | 0.83+/-0.13 | 0.59+/-0.18 | 0.85+/-0.10 | t(65)=0.67 | NS | t(21)=5.37 | *** |
| >3mm | 0.00+/-0.00 | 0.00+/-0.00 | 0.00+/-0.01 | t(27)=2.79 | ** | t(27)=2.76 | * |

## Discussion

Head motion in adults and neonates was found to be different in magnitude, spatial modes and distribution through time. Much of the time, infants actually moved less than adults, but if they moved, the extent was substantially larger. To increase the rate of success in clinical or scientific neonatal imaging sessions, it will be important to mitigate the effects of motion. Some suggestions are discussed below.

Infants showed a greater number of abrupt, large, and multi-axis movements compared to adults, as illustrated in the examples (Figure 1) and quantified by the analysis of the movement magnitude (Figure 4, Table 1). This will make prospective motion correction, in which motion is measured and scanning adjusted, less effective in infants than adults, as faster and more rapid correction will be required. This will be particularly true for methods that have a longer intrinsic lag in the application of correction, such as fMRI (i.e., navigator) correction methods (Thesen et al., 2000) in which motion is estimated from EPI volume and after analysis, corrected in subsequent volumes (*~* 2-4 seconds later). Prospective motion correction appears much better suited to adult movement tracks, which involve a more gradual drift in position (Figure 1), and infant researchers should perhaps wait until robust correction has been proven in adults.

The anisotropy of motion in both adults and infants can be used to inform the choice of MRI acquisitions (Figures 2 and 3). In both groups, there is relatively less motion, for example, in *x* translation. For 2D acquisitions that are to be visually inspected, or where slice-by-slice motion correction is to be performed [28], in-plane motion is less disruptive than through-plane motion, so sagittal acquisitions may be more effective.

Importantly, we have shown that it is not just the magnitude of motion that can differ between groups, but also the spatial patterns of motion; furthermore, it is feasible that RMS motion might be similar between groups, yet they will differ in *how* they move. Differences in the spatial modes of motion will likely lead to different spatial patterns of sensitivity in functional connectivity MRI across groups. Special care must be taken, therefore, when concluding that patterns of connectivity differ between groups, to ensure that this does not just reflect differential motion artifacts. Similar consideration should be given to the potential for different spatial inhomogeneities in sensitivity in anatomical and diffusion-weighted imaging when comparing these across groups.

The more abrupt motion in infants will make it important to evaluate the choice of MRI protocol specifically in this group. Furthermore, the different spatial modes will differently affect different sequences and protocols, and so the rank ordering of signal-to-noise across protocols may change between the groups. We have recently found that multiband EPI sequences to be more robust to motion than standard EPI, with a peak at around an acceleration factor of 4 (Linke et al, submitted). Similarly, the differences in the magnitude and pattern of movement may change which motion correction and denoising strategies are most effective during postprocessing, and it will be important to conduct evaluations of these specifically in infant MRI data. We have found that slice-by-slice realignment is robust when properly regularlized, and increases SNR in infant fMRI (Wild et al, in preparation).

Although we used a gradient-echo EPI sequence (typically used for fMRI) to measure motion, the results are applicable to other kinds of MRI sequences disrupted by motion, such as Diffusion Weighted Imaging or structural (e.g., T1, T2) imaging.

A limitation of our study is that neonatal movement was measured in a particular patient group. In other work (not reported here) we have found that infants younger than 3 months sleep more readily in the MRI scanner, and as a result move less than older ones. It is likely that our patient group, due to premature birth or other injury, lag behind in development. Furthermore, it is likely that as NICU inpatients, they were frailer than typically developing infants. Thus, typically developing infants may move more. Although our measurements are (to our knowledge) the best available characterization of neonatal movement, care should be taken when generalizing to the design of studies of typically developing infants.

Another limitation is that different field strengths were used to measure movement in infants and adults. However, image quality was good on both MRI scanners, and estimating motion involves a relatively small number of parameters (6) from approximately 100 000 data points, and so is robust. We are aware of no evidence showing that the differences in signal-to-noise ratio between 1.5 and 3 T scanners meaningfully impact the precision of motion estimation.

There are a number of potential extensions to this work. An important and pressing extension is to other age and patient groups. It would also be useful to formally evaluate what other factors affect motion, such as: the stimulation paradigm or task; the arousal state of the patient; and the acoustic noise of specific MRI sequences (by measuring movement in an MRI independent way such as by an ultrasonic or optical movement tracker).

In sum, movement in neonates was found to differ from adults substantially in magnitude and form, and this should be accounted for when designing MRI acquisitions and analyses in this group. More generally, this study forms a proof-of-principle that motion should be measured for different patient groups, and acquisition and analysis choices adjusted accordingly.

## Acknowledgements

We would like to thank Linda Geerligs, for a useful conversation during the genesis of this project, and Chao-Gan Yan for providing open source matlab scripts to calculate root-mean square displacement. For funding, we thank NSERC/CIHR CHRP (201110CPG), NSERC Discovery and the Canada Excellence Research Chair (CERC) in Cognitive Neuroimaging.

